# Metacognition and Problem Solving: How Self-Coaching Helps First-Year Students Move Past the Discomfort of Monitoring

**DOI:** 10.1101/2023.08.16.553589

**Authors:** Stephanie M. Halmo, Kira A. Yamini, Julie Dangremond Stanton

## Abstract

Stronger metacognitive regulation skills are linked to increased academic achievement. Metacognition has primarily been studied using retrospective methods, but these methods limit access to students’ in-the-moment metacognition. We investigated first-year life science students’ in-the-moment metacognition while they solved challenging problems, and asked 1) What metacognitive regulation skills are evident when first-year life science students solve problems on their own? and 2) What aspects of learning self-efficacy do first-year life science students reveal when they solve problems on their own? Think aloud interviews were conducted with 52 first-year life science students across three institutions and analyzed using qualitative content analysis. Our results reveal that first-year life science students use an array of monitoring and evaluating skills while solving problems, which challenges the deficit-oriented notion that students enter college with poor metacognitive skills. Additionally, a handful of students self-coached or encouraged themselves as they confronted aspects of the problems that were unfamiliar. These verbalizations suggest ways we can encourage students to couple their metacognitive regulation skills and self-efficacy to persist when faced with challenging disciplinary problems. Based on our findings, we offer recommendations for how instructors can help first-year life science students develop and strengthen their metacognition to achieve improved problem-solving performance.

## INTRODUCTION

Have you ever asked a student to solve a problem, seen their solution, and then wondered what they were thinking while they were problem solving? As college instructors, we often ask students in our classes to solve problems. Sometimes we gain access to our student’s thought process or cognition through strategic question design and direct prompting. Far less often we gain access to how our students regulate and control their own thinking, or the metacognition they use in the moment to solve. Retrospective methods can and have been used to access this information from students, but students often cannot remember what they were thinking a week or two later. We lack deep insight into students’ in-the-moment metacognition because it is challenging to obtain their in-the-moment thoughts. Not having access to students’ metacognition in-the-moment presents a barrier towards developing effective metacognitive interventions to improve learning. Educators and students alike are interested in metacognition because of its malleable nature and demonstrated potential to improve academic performance. Metacognition of life science undergraduates has been studied widely using retrospective methods. In contrast, less is known about how life science undergraduates use their metacognition through in-the-moment methods like think aloud interviews. Thus, there is a need to characterize how life science undergraduates use their metacognition during individual problem-solving and to offer evidence-based suggestions to instructors for supporting students’ metacognition. In particular, understanding the metacognitive skills first-year life science students bring to their introductory courses will position us to better support their learning earlier on in their college careers and set them up for future academic success.

### Metacognition

Metacognition, or one’s awareness and control of their own thinking for the purpose of learning (Cross & Paris, 1988), is linked to improved academic achievement. In one meta-analysis of studies that spanned developmental stages from elementary school to adulthood, metacognition predicted academic performance when controlling for intelligence (Ohtani & Hisasaka, 2018). In another meta-analysis specific to mathematics, researchers found a significant positive correlation between metacognition and math performance in adolescences, indicating individuals who demonstrated stronger metacognition also performed better on math tasks (Muncer et al., 2022). The strong connection between metacognition and academic achievement represents a potential leverage point for enhancing student learning and success in the life sciences. If we explicitly teach life science undergraduates how to develop and use their metacognition, we can expect to increase the effectiveness of their learning and subsequent academic success. However, in order to provide appropriate guidance, we must first know how students in the target population are employing their metacognition.

Based on one theoretical framework of metacognition, metacognition is comprised of two components: metacognitive awareness and metacognitive regulation (Schraw & Moshman, 1995). Metacognitive awareness includes one’s knowledge of learning strategies and of themselves as a learner. Metacognitive regulation encompasses how students act on their metacognitive awareness or the actions they take to learn (Sandi-Urena et al., 2011). Metacognitive regulation is broken up into three skills: *planning* how to approach a learning task or goal, *monitoring* progress towards achieving that learning task or goal, and *evaluating* achievement of said learning task or goal (Stanton et al., 2021). These regulation skills can be thought of temporally: planning occurs before learning starts, monitoring occurs during learning, and evaluating takes place after learning has occurred. As biology education researchers, we are particularly interested in life science undergraduates’ metacognitive regulation skills or the actions they take to learn because regulation skills have been shown to have a more dramatic impact on learning than awareness alone (Dye & Stanton, 2017).

Importantly, metacognition is context-dependent, meaning metacognition use may vary depending on factors such as the subject matter or learning task (Kelemen et al., 2000; Kuhn, 2000; Veenman & Spaans, 2005). For example, the metacognitive regulation skills a student may use to evaluate their learning after reading a text in their literature course may differ from those skills the same student uses to evaluate their learning on a genetics exam. This is why it is imperative to study metacognition in a particular context, like the life sciences. Metacognition is often thought of as a domain-general skill because of its broad applicability across different disciplines. However, metacognitive skills are first developed in a very domain-specific way and then those metacognitive skills can become more generalized over time as they are further developed and honed (Kuhn, 2000; Veenman & Spaans, 2005). This is in alignment with research from the problem-solving literature that suggests stronger problem-solving skills are a result of deep knowledge within a domain (Frey et al., 2022; Pressley et al., 1987). For example, experts are known to classify problems based on deep conceptual features because of their well-developed knowledge base whereas novices tend to classify problems based on superficial features (Chi et al., 1981).

### Methods for Studying Metacognition

Researchers use two main methods to study metacognition: retrospective and in-the-moment methods. Retrospective methods ask learners to reflect on learning they’ve done in the past. In contrast, in-the-moment methods ask learners to reflect on learning they’re currently undertaking (Veenman et al., 2006). Retrospective methods include self-report data from surveys like the Metacognitive Awareness Inventory (Schraw & Dennison, 1994) or exam “wrappers” or self-evaluations (Hodges et al., 2020). Whereas in-the-moment methods include think-aloud interviews, which ask students to verbalize all of their thoughts while they solve problems (Bannert & Mengelkamp, 2008; Blackford et al., 2023; Ku & Ho, 2010), or online computer chat log-files as groups of students work together to solve problems (Hurme et al., 2006; Zheng et al., 2019).

Most metacognition research on life science undergraduates, including our own work, has utilized retrospective methods (Dye & Stanton, 2017; Stanton et al., 2019; Stanton et al., 2015). Important information about first-year life science students’ metacognition has been gleaned using retrospective methods, particularly in regard to planning and evaluating. For example, first-year life science students tend to use strategies that worked for them in high school, even if they do not work for them in college, suggesting first-year life science students may have trouble evaluating their study plans (Stanton et al., 2015). Additionally, first-year life science students abandon strategies they deem ineffective rather than modifying them for improvement (Stanton et al., 2019). Lastly, first-year life science students are willing to change their approach to learning, but they may lack knowledge about which approaches are effective or evidence-based (Stanton et al., 2015; Tomanek & Montplaisir, 2004).

In both of the meta-analyses described at the start of this *Introduction*, the effect sizes were larger for studies that used in-the-moment methods (Muncer et al., 2022; Ohtani & Hisasaka, 2018). This means the predictive power of metacognition for academic performance was more profound for studies that used in-the-moment methods to measure metacognition compared to studies that used retrospective methods. One implication of this finding is that studies using retrospective methods might be failing to capture metacognition’s profound effects on learning and performance. Less research has been done using in-the-moment methods to study metacognition in life science undergraduates likely because of the time-intensive nature of collecting and analyzing data using these methods. One study that used think-aloud methods to investigate biochemistry students’ metacognition when solving open-ended buffer problems found that monitoring was the most commonly used metacognitive regulation skill (Heidbrink & Weinrich, 2021). Another study that used think-aloud methods to explore Dutch third-year medical school students’ metacognition when solving physiology problems about blood flow also revealed a focus on monitoring, with students also planning and evaluating but to a lesser extent (Versteeg et al., 2021). Further investigation into the nature of the metacognition first-year life science students use when solving problems is needed in order to provide guidance to this population and their instructors on how to effectively use and develop their metacognitive regulation skills.

### Metacognition and Self-efficacy

Metacognition is related to self-efficacy, another construct that impacts learning. Self-efficacy is one’s belief in their capability to carry out a task (Bandura, 1977; Bandura, 1997). Research on self-efficacy has revealed its predictive power in regards to performance, academic achievement, and selection of a college major (Pajares, 1996). The large body of research on self-efficacy suggests that students who believe they are capable academically, engage more metacognitive strategies, and persist to obtain academic achievement compared to those who do not. In STEM in particular, studies tend to reveal gender differences in self-efficacy with undergraduate men indicating higher self-efficacy in STEM disciplines compared to women (Stewart et al., 2020). In one study of first-year biology students, women were significantly less confident than men and students’ biology self-efficacy increased over the course of a single semester when measured at the beginning and end of the course (Ainscough et al., 2016). However, self-efficacy is known to be a dynamic construct, meaning one’s perception of their capability to carry out a task can vary widely across different task types and over time as struggles are encountered and expertise builds for certain tasks (Yeo & Neal, 2006).

Both metacognition and self-efficacy are strong predictors of academic achievement and performance. For example, one study found that students with stronger metacognitive regulation skills and greater self-efficacy beliefs (as measured by self-reported survey responses) perform better and attain greater academic success (as measured by GPA) (Coutinho & Neuman, 2008). Additionally, self-efficacy beliefs were strong predictors of metacognition, suggesting students with higher self-efficacy used more metacognition. Together, the results from this quantitative study using structural equation modeling of self-reported survey responses suggests that metacognition may act as a mediator in the relationship between self-efficacy and academic achievement (Coutinho & Neuman, 2008). As qualitative researchers, we were curious to uncover how both metacognition and self-efficacy might emerge out of more qualitative, in-the-moment data streams.

### Research Questions

To pinpoint first-year life science students’ metacognition in-the-moment and to describe the relationship between their metacognition and self-efficacy, we conducted think aloud interviews with 52 students from three different institutions to answer the following research questions:

1. What metacognitive regulation skills are evident when first-year life science students solve problems on their own?
2. What aspects of learning self-efficacy do first-year life science students reveal when they solve problems on their own?

## METHODS

### Research Participants & Context

This study is a part of a larger longitudinal research project investigating the development of metacognition in life science undergraduates which was classified by the Institutional Review Board at the University of Georgia (STUDY00006457) and University of North Georgia (2021-003) as exempt. For that project, 52 first-year students at three different institutions in the southeastern United States were recruited from their introductory biology or environmental science courses in the 2021-2022 academic year. Data was collected at three institutions (Georgia Gwinnett College, University of Georgia, and University of North Georgia) to represent different academic environments because it is known that context can affect metacognition (see **Supplemental Data Table 1**). Additionally, in our past work we found that first-year students from different institutions differed in their metacognitive skills (Stanton et al., 2019; Stanton et al., 2015). Our goal in collecting data from three different institution types was to ensure our qualitative study could be more generalizable than if we had only collected data from one institution. Students at each institution were invited to complete a survey to provide their contact information, answer the revised 18-item Metacognitive Awareness Inventory (Harrison & Vallin, 2018), 32-item Epistemic Beliefs Inventory (Schraw et al., 1995), and 8-item Self-efficacy for Learning and Performance subscale from the Motivated Strategies for Learning Questionnaire (MSLQ) (Pintrich et al., 1993). They were also asked to provide their demographic information including their age, gender, race/ethnicity, college experience, intended major, and first-generation status. First-year students who were 18 years or older and majoring in the life sciences were invited to participate in the larger study. We used purposeful sampling to select a sample that matched the demographics of the student body at each institution and also represented a range in metacognitive ability based on students’ responses to the revised Metacognitive Awareness Inventory (Harrison & Vallin, 2018). In total, eight students from Georgia Gwinnett College, 23 students from the University of Georgia, and 21 students from the University of North Georgia participated in the present study (see **Supplemental Data Table 2**).

### Data Collection

All interviews were conducted over Zoom during the 2021-2022 academic year when participants had returned to the classroom. Participants (n=52) were asked to think aloud as they solved two challenging biochemistry problems (**Figure 1**) that have been previously published (Bhatia et al., 2022; Halmo et al., 2018; Halmo et al., 2020). We selected two challenging biochemistry problems for first-year students to solve because we know that students do not use metacognition unless they find a learning task challenging (Carr & Taasoobshirazi, 2008). The problems we selected met this criterion because participants had not yet taken biochemistry. The biology problems were open-ended and asked students to make predictions and provide scientific explanations for their predictions about 1) non-covalent interactions in a folded protein for the Protein X Problem (Halmo et al., 2018; Halmo et al., 2020) and 2) negative feedback regulation in a metabolic pathway for the Pathway Flux Problem (Bhatia et al., 2022). To elicit student thinking after participants fell silent for more than five seconds, interviewers used the following two prompts: “What are you thinking (now)?” and “Can you tell me more about that?” (Charters, 2003; Ericsson & Simon, 1980). After participants solved the problems, they shared their written solutions with the interviewer using the chat feature in Zoom. Participants were then asked to describe their problem-solving process out loud and respond to up to four reflection questions (see **Supplemental Material** for full interview protocol). The think aloud interviews were audio and video recorded and transcribed using a professional, machine-generated transcription service (Temi.com). All transcripts were checked for accuracy by members of the research team before analysis began.

**Figure 1.**
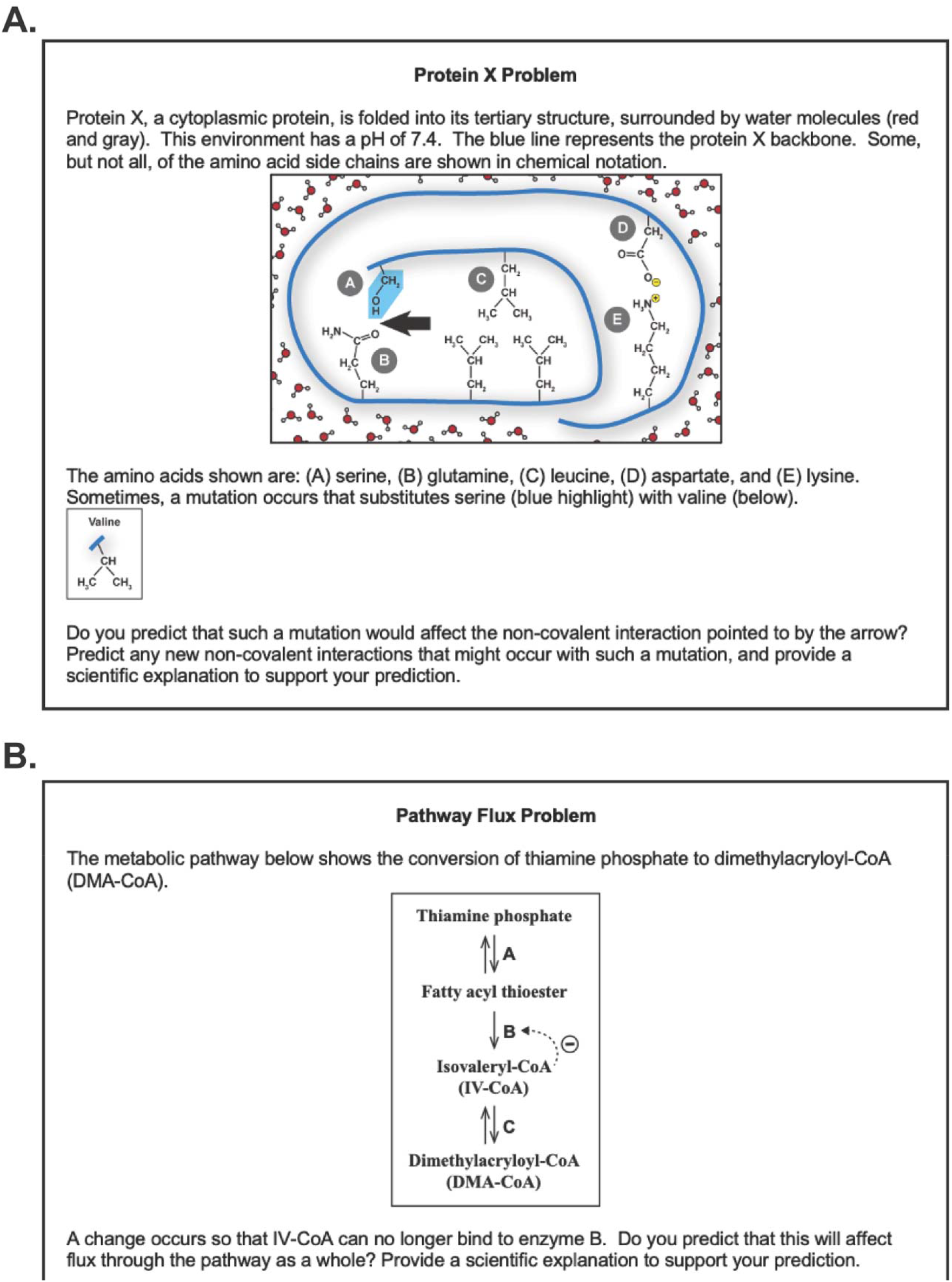
Think Aloud Problems. Students were asked to think aloud as they solved two challenging biochemistry problems. Panel A depicts the Protein X Problem previously published in Halmo et al., 2018 and Halmo et al., 2020. Panel B depicts the Pathway Flux problem previously published in Bhatia et al., 2022. Both problems are open-ended and ask students to make predictions and provide scientific explanations for their predictions.

### Data Analysis

The resulting transcripts were analyzed by a team of three researchers in three cycles. In the first cycle of data analysis, half of the transcripts were open coded by members of the research team (S.M.H., J.D.S., and K.A.Y.). During this open coding process, we individually reflected on the contents of the data, remained open to possible directions suggested by our interpretation of the data, and recorded our initial observations using analytic memos (Saldaña, 2021). The research team (S.M.H., J.D.S., and K.A.Y.) then met to discuss our observations from the open coding process and suggest possible codes that were aligned with our observations, knowledge of metacognition and self-efficacy, and our guiding research questions. This discussion led to the development of an initial codebook consisting of inductive codes discerned from the data and deductive codes derived from theory on metacognition. In the second cycle of data analysis, the codebook was applied to the dataset iteratively by two researchers (S.M.H. and K.A.Y) using MaxQDA2020 software (VERBI Software; Berlin, Germany) until the codebook stabilized. Coding disagreements between the two coders were discussed by all three researchers until consensus was reached. All transcripts were coded to consensus to identify aspects of metacognition and learning self-efficacy that were verbalized by participants. Coding to consensus allowed the team to consider and discuss their diverse interpretations of the data and ensure trustworthiness of the analytic process (Tracy, 2010). In the third and final cycle of analysis, thematic analysis was used to uncover central themes in our dataset. As a part of thematic analysis, two researchers (S.M.H and K.A.Y) synthesized one-sentence summaries of each participant’s think aloud interview. Student quotes presented in the Results & Discussion have been lightly edited for clarity, and all names are pseudonyms.

### Problem-Solving Performance as Context for Studying Metacognition

Participants’ final problem solutions were individually scored by two researchers (S.M.H. and K.A.Y) using an established rubric and scores were discussed until complete consensus was reached. The median problem-solving performance of students in our sample was two points on a 10-point rubric. Students in our sample scored low on the rubric because they either failed to answer part of the problem or struggled to provide accurate explanations or evidence to support their predictions. Despite the phrase “provide a scientific explanation to support your prediction” included in the prompt, most students’ solutions contained a prediction, but lacked an explanation. For example, the majority of the solutions for the Protein X problem predicted the non-covalent interaction would be affected by the substitution, but lacked categorization of the relevant amino acids or identification of the non-covalent interactions involved, which are critical problem-solving steps for this problem (Halmo et al., 2018; Halmo et al., 2020). The majority of the Pathway Flux solutions also predicted that flux would be affected, but lacked an accurate description of negative feedback inhibition or regulation release of the pathway, which are critical features of this problem (Bhatia et al., 2022). This lack of accurate explanations is not unexpected. Previous work shows that both introductory biology and biochemistry students struggle to provide accurate explanations to these problems without pedagogical support, and introductory biology students generally struggle more than biochemistry students (Bhatia et al., 2022; Lemons, personal communication).

The problem-solving scores were interrogated using R Statistical Software (R Core Team, 2021). A one-way ANOVA was performed to compare the effect of institution on problem-solving performance. This analysis revealed that there was not a statistically significant difference in problem-solving performance between the three institutions (F(2, 49) = 0.085, p = 0.92). This indicates students performed similarly on the problems regardless of which institution they attended (**Supplemental Data Table 3**). Another one-way ANOVA was performed to compare the effect of gender on problem-solving performance which revealed no statistically significant differences in problem-solving performance based on gender (F(1, 50) = 0.956, p = 0.33). Students performed similarly on the problems regardless of their gender (**Supplemental Data Table 4**). Taken together, this analysis suggests a homogeneous sample in regard to problem-solving performance.

## RESULTS & DISCUSSION

### What metacognitive regulation skills are evident when first-year life science students solve problems on their own?

To address our first research question, we looked for statements and questions related to the three skills of planning, monitoring, and evaluating in our participants’ think aloud data. Because metacognitive regulation skills encompass how students act on their metacognitive awareness, participants’ explicit awareness was a required aspect when analyzing our data for these skills. For example, the statement “this is a hydrogen bond” does not display awareness of one’s knowledge but rather the knowledge itself (cognition). In contrast, the statement “I know this is a hydrogen bond” does display awareness of one’s knowledge and is therefore considered evidence of metacognition. We present our findings for each metacognitive regulation skill. For further demonstration of how students use these skills in concert when problem solving, we offer problem-solving vignettes of a student from each institution in **Supplemental Data**.

#### Planning: Students did not plan before solving but did assess the task and rationalize their approach in the moment

Planning how to approach the task of solving problems individually involves selecting strategies to use and when to use them before starting the task (Stanton et al., 2021). Planning did not appear in our data in a classical sense. This finding is unsurprising because the task was 1) well-defined, meaning there were a few potentially accurate solutions rather than an abundant number of accurate solutions, 2) straightforward meaning the goal of solving the problem was clearly stated, and 3) relatively short meaning students were not entering and exiting from the task multiple times like they might when studying for an exam. Additionally, the stakes were comparatively low meaning task completion and performance carried little to no weight in participants’ college career. In other data from this same sample, we know that these participants make plans for high-stakes assessments like exams but often admit to not planning for lower stakes assessments like homework (unpublished data; Stanton, personal communication). While planning was absent in the traditional sense, it was present in different ways that could also be categorized as planning. Related to the skill of planning, we observed students assessing the task after reading the problem and describing their rationales for their approach in the moment (**Table 1**). We describe these aspects related to planning and provide descriptions of what happened after students planned in this way.

**Table 1.**
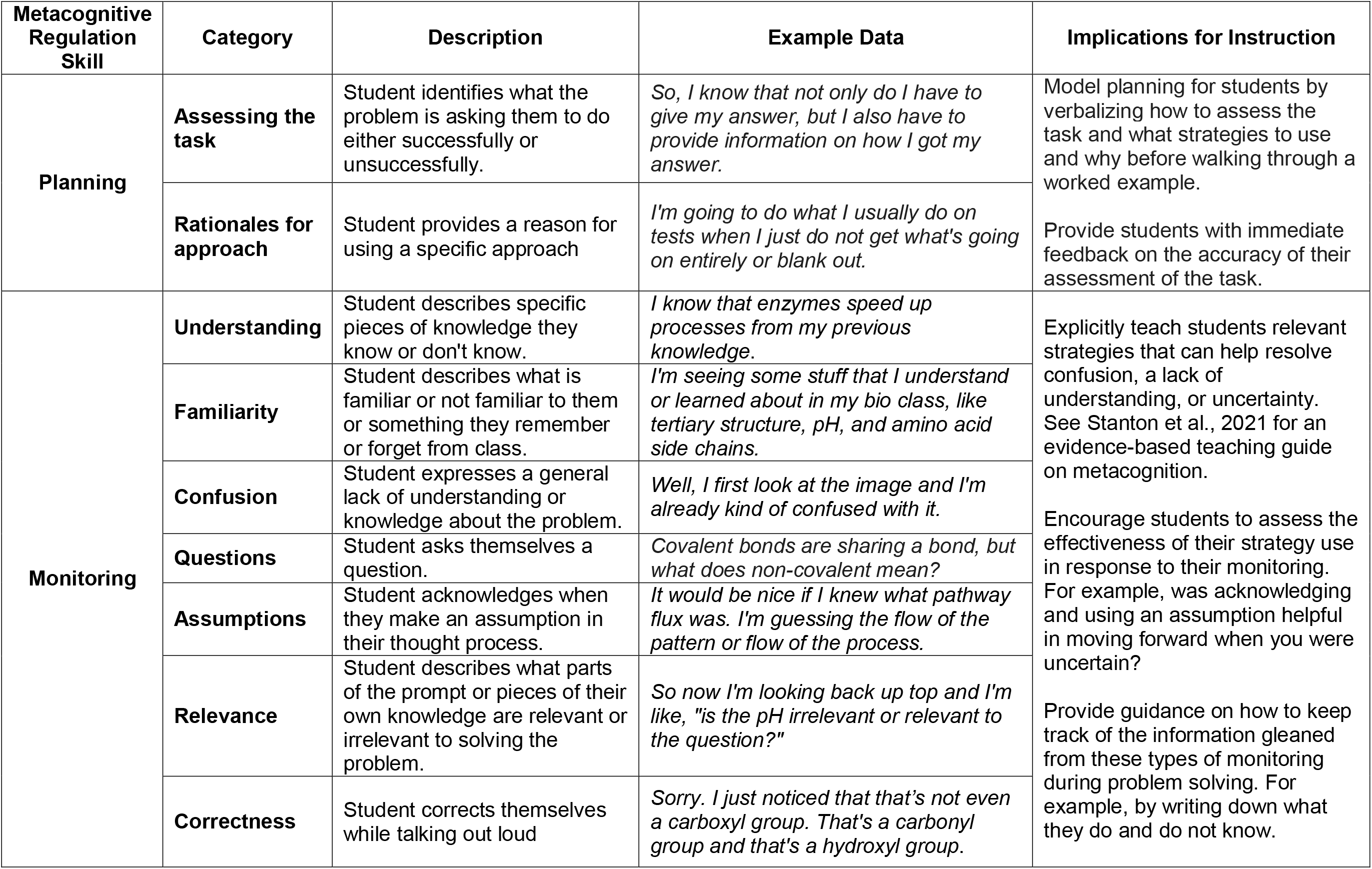
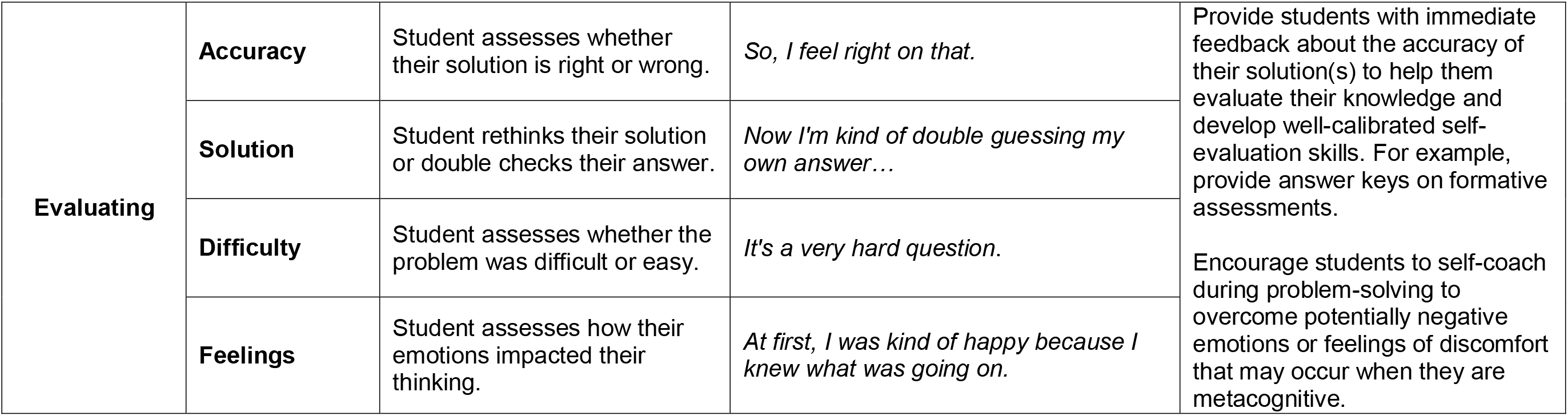
Metacognitive Regulation Skills Revealed during Individual Problem Solving & Implications for Instruction.

##### Assessing the task

While we did not observe students explicitly planning their approach to problem solving *before* beginning the task, we did observe students assessing the task or what other researchers have called “orientation” *after* reading the problems (Meijer et al., 2006; Schellings et al., 2013). Students in our study either assessed the task successfully or unsuccessfully. For example, when Gerald states, *“So I know that not only do **I have to give my answer, but I also have to provide information on how I got my answer**…”* he successfully identified what the problem was asking him to do by providing a scientific explanation. In contrast, Simone admits her struggle with figuring out what the problem is asking when she states, *“I’m still trying to figure out what the question’s asking. I don’t want to give up on this question just yet, but yeah, it’s just kinda hard because **I can’t figure out what the question is asking me** if I don’t know the terminology behind it.”* In Simone’s case, the terminology she struggled to understand is what was meant by a scientific explanation. Assessing the task unsuccessfully also involved misinterpreting what the problem asked. This was a frequent issue for students in our sample during the Pathway Flux problem because students inaccurately interpreted the negative feedback loop, which is a known problematic visual representation in biochemistry (Bhatia et al., 2022). For example, students like Paulina and Kathleen misinterpreted the negative feedback loop as enzyme B no longer functioning when they stated respectively, *“So if enzyme B is taken out of the graph…”*, or *“…if B cannot catalyze…”* Additionally some students misinterpreted the negative feedback loop as a visual cue of the change described in the problem prompt (IV-CoA can no longer bind to enzyme B). This can be seen in the following example quote from Mila: *“So I was looking at it and I see what they’re talking about with the IV-CoA no longer binding to enzyme B and I think that’s what that arrow with the circle and the line through it is representing. It’s just telling me that it’s not binding to enzyme B.”*

###### What happened after assessing the task?

Misinterpretations of what the problem was asking like those shared above from Simone, Paulina, Kathleen, and Mila led to inaccurate answers for the Pathway Flux problem. In contrast, when students like Gerald could correctly interpret what the problem asked them to do, this led to more full and accurate answers for both problems. Accurately interpreting what a problem is asking you to do is critical for problem-solving success. A related procedural error identified in other research on written think aloud protocols from students solving multiple-choice biology problems was categorized as misreading (Prevost & Lemons, 2016).

##### Rationales for approach

We also observed students explaining their rationale for their approach during and after solving. Even though their rationales were not revealed before they started, we still consider their rationales for their approach to be a part of planning because they include their reasoning behind selecting certain strategies. Overall, students revealed rationales for their approach based on their past experience solving problems. Students used two approaches based on their past experience solving exam problems. The first was using instructor recommended strategies for writing a solution. While typing her solution, Elena described, *“I like to like answer the full question, like repeat the question in a short form to answer the question just so it seems like a formal answer. I don’t know. **That’s what my AP biology teacher told me to do**.”* It’s notable that students are remembering strategies their past instructors taught them because it suggests some students are open to recommendations from their current instructors. The second approach students used based on past experience was reading the question first and skipping over the full prompt. As Erwin shared, *“Well, I usually read what it is asking me to answer first. And so I read the bottom paragraph usually first, because sometimes some information can be misleading in test questions.”* In his reflection after solving, Erwin revealed that this *“habit of reading what it’s asking for first”* is *“**a strategy that I did for the ACT and SAT**, where I could read a question and pretty fast. And so I already jumped the gun. I didn’t really care about the first two paragraphs. And I just focus on the question at hand…”* He further explains in his problem reflection that he uses the rest of the problem in the first two paragraphs *“as a reference to answering a question or making me sound smarter in a question, **so that the professor can give me points**.”* Through these snapshots of his think aloud interview, Erwin is revealing his approach for the think aloud problems was similar to his approach for taking exams. While his approach is potentially beneficial on timed, multiple-choice exams like the ACT and SAT, this approach was likely not as effective when solving untimed, free response problems like the ones in this study.

###### What happened after providing a rationale for approach?

When students provided a rationale for their approach, they followed through with using the approach, independent of whether the approach was ultimately beneficial for problem solving or not. For example, Elena’s use of instructor recommended strategies for writing a solution ensured she attempted answering the problem fully. In contrast, while Erwin’s shortcut of reading the question first may have helped him reduce extraneous cognitive (Paas et al., 2003) or metacognitive load (Valcke, 2002; Wirth et al., 2020), it ultimately caused him to miss critical parts of the problem which he only realized during his problem-solving reflection. In Erwin’s case, his approach may have been the misapplication of a well-known general problem-solving approach of working backwards by beginning with the problem goal (Chi & Glaser, 1985).

##### Implications for Instruction & Research about Planning

In our study, evidence of planning was limited. This suggests that first-year students’ approaches were either unplanned or automatic (Samuels et al., 2005). As metacognition researchers and instructors, we find first-year life science students’ absence of planning *before solving* and presence of assessing the task and rationalizing their approach *during and after solving* illuminating. This means planning is likely one area in which we can help first-year life science students grow their metacognitive skills through practice. While we do not anticipate that undergraduate students will be able to plan how to solve a problem that is unfamiliar to them before reading a problem, we do think we can help students develop their planning skills through modeling when solving disciplinary problems. When modeling problem solving for students, we could make our planning explicit for students by verbalizing how we assess the task and what strategies we plan to use and why. From the problem-solving literature, it is known that experts assess a task by recognizing the deep structure or problem type and what is being asked of them (Chi et al., 1981; Smith et al., 2013). This likely happens rapidly and automatically for experts through the identification of visual and key word cues. Forcing ourselves to think about what these cues might be and alerting students to them through modeling may help students more rapidly develop expert-level schema, approaches, and planning skills. Providing students with feedback on their assessment of a task and whether or not they misunderstood the problem also seems to be critical for problem-solving success (Prevost & Lemons, 2016).

Additionally, our data show that when students have a rationale for an approach, they are likely to follow through with implementing that approach. We can use this to our benefit by encouraging our students to assess the effectiveness of the approach they employ which will help them further develop their metacognitive regulation skills. As students develop stronger rationales for their approach, they may be more likely to use those approaches. For instance, helping students realize they can plan for smaller tasks like solving a problem by listing the pros and cons of relevant strategies and what order they plan to use selected strategies before they begin could help students narrow the problem solving space, approach the task with focus, and achieve efficiency to become “good strategy users” (Pressley et al., 1987).

#### Monitoring: Students monitored their knowledge in the moment in a myriad of ways

Monitoring progress towards problem-solving involves assessing conceptual understanding during the task (Stanton et al., 2021). First-year life science students in our study monitored their conceptual understanding during individual problem solving in a myriad of ways. In our analysis, we captured the specific aspects of conceptual understanding students monitored. Students in our sample monitored their 1) understanding, 2) familiarity, 3) confusion, 4) questions, 5) assumptions, 6) relevance, and 7) correctness (**Table 1**). We describe each aspect of conceptual understanding that students monitored and we provide descriptions of what happened after students monitored in this way.

##### Monitoring Understanding

When students monitored understanding, they described specific pieces of knowledge they either knew or did not know, beyond what was provided in the problem prompt. For example, Kathleen demonstrated an awareness of her understanding about amino acid properties when she said, *“**I know** that like the different amino acids all have different properties like some are, what’s it called? Like hydrophobic, hydrophilic, and then some are much more reactive.”* Willibald monitored his understanding of the turn of phrase “when in doubt, van der Waals it out” by sharing, *“So, cause **I know** basically everything has, well not basically everything, but a lot of things have van der Waal forces in them. So that’s why I say that a lot of times. But it’s a temporary dipole, I think.”* In contrast, Jeffery monitored his lack of understanding of a specific part of the Pathway Flux figure when he stated, *“I guess **I don’t understand** what this dotted arrow is meaning.”* Ignoring or misinterpreting the negative feedback loop was a common issue as students solved this problem, so it’s notable that Jeffery acknowledged his lack of understanding about this symbol. When students identified what they knew, the knowledge they revealed sometimes had the potential to lead to a misunderstanding. Take for example Lucy’s quote: *“**I know** a hydrogen bond has to have a hydrogen. I know that much. And it looks like they both have hydrogen.”* This statement suggests Lucy might be displaying a known misconception about hydrogen bonding – that all hydrogens participate in hydrogen bonding (Villafañe et al., 2011).

###### What happened after monitoring understanding?

When students could identify what they knew, they used this information to formulate a solution. When students could identify what they did not know, they either did not know what to do next or they used strategies to move beyond their lack of understanding. Two strategies students used after identifying a lack of understanding included disregarding information and writing what they knew. Kyle disregarded information when he didn’t understand the negative feedback loop in the Pathway Flux problem: *“…there is another arrow on the side I see with a little minus sign. I’m not sure what that means… it’s not the same as [the arrows by] A and C. So I’m just going to disregard it sort of for now. It’s not the same. Just like note that in my mind that it’s not the same.”* In this example, Kyle disregards a critical part of the problem, the negative feedback loop, and does not revisit the disregarded information which ultimately led him to an incorrect prediction for this problem. We also saw one example of a student, Elaine, use the strategy of writing what she knew when she was struggling to provide an explanation for her answer: *“I should know this more, but I don’t know, like a specific scientific explanation answer, but I’m just going to write what I do know so I can try to organize my thoughts.”* Elaine’s focus on writing what she knew allowed her to organize the knowledge she did have into a plausible solution that specified which amino acids would participate in new non-covalent interactions *(“I predict there will be a bond between A and B and possibly A and C.”*) despite not knowing *“what would be required in order for it to create a new noncovalent interaction with another amino acid.”* The strategies that Kyle and Elaine used in response to their monitoring of a lack of understanding shared the common goal of helping them get unstuck in their problem-solving process.

##### Monitoring Familiarity

When students monitored familiarity, they described knowledge or aspects of the problem prompt that were familiar or not familiar to them. This category also captured when students would describe remembering or forgetting something from class. For example, when Simone states, *“**I remember** learning covalent bonds in chemistry, but **I don’t remember** right now what that meant”* she is acknowledging her familiarity with the term covalent from her chemistry course. Similarly, Oliver acknowledges his familiarity with tertiary structure from his class when solving the Protein X problem. He first shared*, “**This reminds me of something** that we’ve looked at in class of a tertiary structure. It was shown differently but **I do remember something similar to this**.”* Then later, he acknowledges his lack of familiarity with the term flux when solving the Pathway Fux problem, *“That word flux. **I’ve never heard that word before**.”* Quinn aptly pointed out that being familiar with a term or recognizing a word in the problem did not equate to her understanding, *“I mean, I know amino acids, but that doesn’t… like **I recognize the word**, but it doesn’t really mean anything to me. And then non-covalent, I recognize the conjunction of words, but again, it’s like somewhere deep in there…”*

###### What happened after monitoring familiarity?

When students recognized what was familiar to them in the problem, it sometimes helped them connect to related prior knowledge. In some cases, though, students connected words in the problem that were familiar to them to unrelated prior knowledge. Erika, for example, revealed in her problem reflection that she was familiar with the term mutation in the Protein X problem and formulated her solution based on her knowledge of the different types of DNA mutations, not non-covalent interactions. In this case, Erika’s familiarity with the term mutation and failure to recognize this familiarity when problem solving impeded her development of an accurate solution to the problem. This is why Quinn’s recognition that her familiarity with terms does not equate to understanding is critical. This recognition can help students like Erika avoid false feelings of knowing that might come from the rapid fluent recall of unrelated knowledge (Reber & Greifeneder, 2017). When students recognized parts of the problem they were unfamiliar with, they often searched for familiar terms to use as footholds. For example, Lucy revealed the following in her problem reflection: *“So first I tried to look at the beginning introduction to see if I knew anything about the topic. Unfortunately, I did not know anything about it. So I just tried to look for any trigger words that I did recognize.”* After stating this, Lucy said she recognized the words protein and tertiary structure and was able to access some prior knowledge about hydrogen bonds for her solution.

##### Monitoring Confusion

Monitoring confusion is distinct from monitoring understanding. When students displayed awareness of a specific piece of knowledge they did not know (e.g., *“I don’t know what these arrows really mean.”* - Mila) this was considered monitoring (a lack of) understanding. In contrast, monitoring confusion was a more general awareness of their overall lack of understanding (e.g., *“Well, I first look at the image and **I’m already kind of confused with it** [laughs].”* - Erwin). When students monitored confusion when solving, they expressed a general lack of understanding or knowledge about the problem. As Sara put it, *“**I have no clue** what I’m looking at.”* Sometimes monitoring confusion came as an acknowledgement of lack of prior knowledge students felt they needed to solve the problem. Take for instance when Ismail states, *“I’ve never really had any prior knowledge on pathway fluxes and like how they work and it obviously **doesn’t make much sense to me**.”* Students also expressed confusion about how to approach the problem, which is related to monitoring one’s procedural knowledge. For example, when Harper stated, *“**I’m not sure how to approach the question**,”* she was monitoring a lack of knowledge about how to begin. Similarly, after reading the problem Tiffani shared, *“**I am not sure how to solve this one** because I’ve actually never done it before…”* Several of the first-year life science students in our study also got stuck with the request to provide a scientific explanation in the problem prompt, as Simone stated, *“**I don’t know how to provide a scientific explanation** for that.”*

###### What happened after monitoring confusion?

When students monitored their confusion, one of two things happened. Rarely, students would give up on solving altogether. In fact, only one individual (Roland) submitted a final solution that read, “*I have no idea*.” More often students persisted despite their confusion. Rereading the problem was a common strategy students in our sample used after identifying general confusion. As Jeffery stated after reading the problem, *“I didn’t really understand that, so I’m gonna read that again.”* After rereading the problems a few times, Jeffery stated, *“Oh, and we have valine here. I didn’t see that before.”* Some students like Valentina revealed their rereading strategy rationale after solving, *“First I just read it a couple of times because I wasn’t really understanding what it was saying.”* After rereading the problem a few times Valentina was able to accurately assess the task by stating *“amino acid (A) turns into valine.”* When solving, some students linked their general lack of prior knowledge or knowledge about how to proceed with an inability solve. As Harper shared, *“I don’t think that I have enough like basis or learning to where I’m able to answer that question.”* Similarly, Tiffani shared, *“I am actually not sure how to solve this. I do not think I can solve this one.”* Despite making these claims of self-doubt in their ability to solve, both Harper and Tiffani monitored in other ways and ultimately came up with a solution beyond a simple, “I don’t know.” In sum, when students acknowledged their confusion in this study, they usually did not stop there. They used their confusion as a jumping off point to further monitor by identifying more specifically what they did not understand or they used a strategy, like re-reading, to resolve their confusion. Persisting despite confusion is likely dependent on motivational factors which were not explored in this study.

##### Monitoring Questions

When students monitored through questions, they asked themselves a question out loud. These questions were either about the problem itself or their own knowledge. An example of monitoring through a question about the problem itself comes from Elaine who asked herself after reading the problem and sharing her initial thoughts, *“**What is this asking me?**”* Elaine’s question helped reorient her to the problem and put herself back on track with answering the question asked. After Edith came to a tentative solution, she asked herself, *“But what about the other information? **How does that pertain to this?**”* which helped her initiate monitoring the relevance of the information provided in the prompt. Students also posed questions to themselves about their own content knowledge. Take for instance Phillip when he asked himself, *“So would non-covalent be ionic bonds or would it be something else? Covalent bonds are sharing a bond, but **what does non-covalent mean?**”* After Phillip asked himself this question, he reread the problem but ultimately acknowledge he was *“not too sure what non-covalent would mean.”*

###### What happened after monitoring questions?

After students posed questions to themselves while solving, they either answered their question or they did not. Students who answered their self-posed questions relied on other forms of monitoring and rereading the prompt to do so. For example, after questioning themselves about their conceptual knowledge, some students acknowledged they did not know the answer to their question by monitoring their understanding. Students who did not answer their self-posed questions moved on without answering their question directly out loud.

##### Monitoring Assumptions

When students monitored assumptions, they explicitly acknowledged when they were making assumptions in their thought process. In the Pathway Flux problem, the majority of students admitted to not knowing what the term flux meant, which lead them to make some assumptions about its meaning. As Elena put it, *“So when it says flux, I think of flow, like the flow of the pathway. **I’m assuming that’s what it’s asking**.”* Henry acknowledges he made the same assumption as Elena, *“It would be nice if I knew what pathway flux was. **I’m guessing** the flow of the pattern or flow of the process.”* Students like Ismail also acknowledged when they made assumptions about associations between words: *“When I see the word ’non-covalent’**, I presume** ionic interactions are the type of interactions that are being in question.”* In one case, Icarus acknowledged a central assumption pertinent to his more sophisticated prediction to the Pathway Flux problem: *“…if IV-CoA can’t bind to enzyme B as a substrate, **assuming part of IV-CoA in excess**, this is what helps enzyme B work.”* Icarus was one of the only students in our sample to get at the idea of relative amounts of metabolites in the Pathway Flux problem.

###### What happened after monitoring assumptions?

Monitoring assumptions allowed students to continue problem solving rather than getting stuck on what they did not know. Intriguingly, other researchers have identified making incorrect assumptions as a procedural error when solving multiple-choice biology problems (Prevost & Lemons, 2016). We posit that this error may occur when there is a failure to monitor the assumptions being made during problem solving. Explicitly acknowledging when as assumption is made might prevent this procedural error from occurring.

##### Monitoring Relevance

When students monitored relevance, they described what pieces of their own knowledge or aspects of the problem prompts were relevant or irrelevant to their thought process. For the Protein X problem, many students monitored the relevance of the provided information about pH. First-year life science students may have focused on this aspect of the problem prompt because pH is a topic often covered in introductory biology classes, which all participants were enrolled in at the time of the study. However, students differentially decided if this information was relevant or irrelevant. Quinn decided this piece of information was relevant: *“The pH of the water surrounding it. **I think it’s important** because otherwise it wouldn’t really be mentioned.”* In contrast, Ignacio decided the same piece of information was irrelevant: *“So **the pH has nothing to do with it**. The water molecules had nothing to do with it as well. So basically everything in that first half, everything in that first thing, right there is basically useless. **So I’m just going to exclude that information out of my thought process** cause the pH has nothing to do with what’s going on right now…”* From an instructional perspective, knowing the pH in the Protein X problem is relevant information for determining the ionization state of acidic and basic amino acids, like amino acids D and E shown in the figure. However, this specific problem asked students to specifically consider amino acids A and B, so Ignacio’s decision that the pH was irrelevant may have helped him focus on more central parts of the problem. In addition to monitoring the relevance of the provided information, sometimes students would monitor the relevance of their own knowledge that they brought to bear on the problem. For example, consider the following quote from Regan: *“I just think that it might be a hydrogen bond, which **has nothing to do with the question**.”* Regan made this statement during her think aloud for the Protein X problem, which is intriguing because the Protein X problem deals solely with non-covalent interactions like hydrogen bonding.

###### What happened after monitoring relevance?

Overall, monitoring relevance helped students narrow their focus during problem solving, but could be misleading if done inaccurately like in Regan’s case.

##### Monitoring Correctness

When students monitored correctness, they corrected their thinking out loud. A prime example of this comes from Kyle’s think aloud, where he corrects his interpretation of the problem not once but twice: *“It said the blue one highlighted is actually a valine, which substituted the serine, so that’s valine right there. And then I’m reading the question. **No, no, no. It’s the other way around.** So serine would substitute the valine and the valine is below…**Oh wait wait, I had it right the first time.** So the blue highlighted is this serine and that’s supposed to be there, but a mutation occurs where the valine gets substituted.”* Kyle first corrects his interpretation of the problem in the wrong direction but corrects himself again to put him on the right track. Icarus also caught himself reading the problem incorrectly by replacing the word non-covalent with the word covalent, which was a common error students made: *“**Oh, wait, I think I read that wrong.** I think I read it wrong. Well, yeah. Then that will affect it. I didn’t read the non-covalent part. I just read covalent.”* Students also corrected their language use during the think aloud interviews, like Edith: *“**since enzyme B is no longer functioning… No, not enzyme B… since IV-CoA is no longer functional** and able to bind to enzyme B, the metabolic pathway is halted.”* Edith’s language use correction, while minor, is worth noting because students in this study often misinterpreted the Pathway Flux problem to read as “enzyme B no longer works”. There were also instances when students corrected their own knowledge that they brought to bear on the problem. This can be seen in the following quote from Tiffani when she says, *“And tertiary structure. It has multiple… **No, no, no. That’s primary structure.** Tertiary structure’s when like the proteins are folded in on each other.”*

###### What happened after monitoring correctness?

When students corrected themselves, this resulted in more accurate interpretations of the problem and thus more accurate solutions. Specifically, monitoring correctness helped students avoid common mistakes when assessing the task which was the case for Kyle, Icarus, and Edith described above. When students do not monitor correctness, incorrect ideas can go unchecked throughout their problem-solving process, leading to more inaccurate solutions. In other research, contradicting and misunderstanding content were two procedural errors students experienced when solving multiple-choice biology problems (Prevost & Lemons, 2016), which could be alleviated through monitoring correctness.

##### Implications for Instruction & Research about Monitoring

Monitoring is the last metacognitive regulation skill to develop, and it develops slowly and well into adulthood (Schraw, 1998). Based on our data, first-year life science students are monitoring in the moment in a myriad of ways. This may suggest that college-aged students have already developed monitoring skills by the time they enter college. This finding has implications for both instruction and research. For instruction, we may need to help our students keep track of and learn what do with the information and insight they glean from their *in-situ* monitoring when solving disciplinary problems. For example, students in our study could readily identify what they did and did not know, but they may struggle to identify ways in which they could potentially resolve their lack of understanding, confusion, or uncertainty or use this insight in expert-like ways when formulating a solution.

As instructors who teach students about metacognition, we can normalize the temporary discomfort monitoring may bring as an integral part of the learning process and model for students what to do after they monitor. For example, when students glean insight from *monitoring familiarity*, we could help them learn how to properly use this information so that they do not equate familiarity with understanding when practicing problem solving on their own. This could help students avoid the fluency fallacy or the false sense that they understand something simply because they recognize it or remember learning about it (Reber & Greifeneder, 2017). The majority of the research on metacognition, including our own, has been conducted using retrospective methods (Dye & Stanton, 2017; Stanton et al., 2019; Stanton et al., 2015). However, retrospective methods may provide little insight into true monitoring skills since these skills are used *during* learning rather than *after* learning has occurred (Schraw & Moshman, 1995; Stanton et al., 2021). More research using in-the-moment methods, which are used widely in the problem-solving literature, are needed to fully understand the rich monitoring skills of life science students and how they may develop over time. The monitoring skills of life science students in both individual and small group settings, and the relationship of monitoring skills across these two settings, warrants further exploration. This seems particularly salient given that questioning and responding to questions seems to be an important aspect of both individual metacognition in the present study and social metacognition in our prior study, which used in-the-moment methods (Halmo et al., 2022).

#### Evaluating: Students evaluated their knowledge and experience problem solving

Evaluating achievement of individual problem solving involves appraising an implemented plan and how it could be improved for future learning after completing the task (Stanton et al., 2021). Students in our sample revealed some of the ways they evaluate during individual problem solving (**Table 1**). They evaluated both their knowledge and their experience of problem solving. When students in our sample evaluated their knowledge, we categorized this as either accuracy or solution. When students in our sample evaluated their experience, we categorized this as either difficulty or feelings.

##### Evaluating Knowledge: Accuracy

Evaluating accuracy occurred when students assessed whether their solution was right or wrong. For example, when Harper says, *“I think I got the answer right”* she is evaluating her accuracy in the affirmative (that her solution is right). Interestingly, Harper’s answer was only partially correct. In contrast, more students evaluated the accuracy of their solution in the negative. For example, when Kyle states, *“I don’t think hydrogen bonding is correct.”* Kyle clarified in his problem reflection, *“I noticed [valine] did have hydrogens and the only non-covalent interaction I know of is probably hydrogen bonding. So I just sort of stuck with that and just said more hydrogen bonding would happen with the same oxygen over there [in glutamine].”* Through this quote, we see that Kyle went with hydrogen bonding as his prediction because it’s the only non-covalent interaction he could recall. However, Kyle accurately evaluated the accuracy of his solution by noting that hydrogen bonding was not the correct answer. Evaluating accuracy in the negative often seemed like hedging or self-doubt. Take for instance Astrid’s quote about her tentative solution, *“I’m using all my previous knowledge to try and put something together, but **it’s probably not right**.”* After making this statement Astrid was able to move forward in her problem solving to come to a solution that was not fully accurate. Regan’s quote that she shared right after submitting her final solution also expressed self-doubt about the accuracy of her solution: *“**The chances of being wrong are 100%**, just like, you know [laughs].”* While all of the above examples of evaluating accuracy occurred spontaneously without prompting, having students describe their thinking process after solving the problems may have been sufficient to prompt them to evaluate the accuracy of their solution. For example, Erwin evaluated the accuracy of his solution in response to a follow-up question about what he already knew or remembered when solving: *“Well, when I see ”covalent”, it pops into my head of covalent bonding and it’s basically telling me that it’s not covalent bonding, so I have to assume it’s like…ah, crap. **I answered it wrong didn’t I? [laughing] The more I say it, the more I realize that I just answered it completely wrong** because I just wrote down what I was thinking, but I guess that’s okay.”* Erwin’s evaluation of accuracy is accurate because in his solution he incorrectly discussed covalent bonding and not non-covalent interactions.

###### What happened after evaluating accuracy?

When students evaluated the accuracy of their solution it helped them recognize potential flaws or mistakes in their answers. Additionally, acknowledging the possibility that their solutions might be wrong seemed to helped some students continue problem solving as was the case for Astrid.

##### Evaluating Knowledge: Solution

The other way students in our study evaluated their knowledge was through evaluating their solution. Evaluating solutions occurred when students would double-check or rethink their solution. Kyle used a very clearly-defined approach for double checking his work by solving the problem twice: *“So that’s just my initial answer I would put, and then what I do next was **I’d just like reread the question and sort of see if I come up with the same answer after rereading and redoing the problem.** So I’m just going to do that real quick.”* Checking one’s work is a well-established problem-solving step that most successful problem solvers undertake (Cartrette & Bodner, 2010; Prevost & Lemons, 2016). Other students also rethought their initial solution, although their evaluations of their solution seemed less planned than Kyle’s. In the following case, Mila’s evaluation of her solution did not improve her final answer. Mila initially predicted that the change described in the Pathway Flux problem would affect flux, which is correct. However, she evaluates her solution when she states, *“Oh, wait a minute, now that I’m saying this out loud, I don’t think it’ll affect it because I think IV-CoA will be binding to enzyme B or C. Sorry, hold on. **Now I’m like rethinking my whole answer**.”* After this evaluation, Mila changes her prediction to *“it won’t affect flux”*, which is incorrect. In contrast, some students’ evaluations of their solutions resulted in improved final answers. For example, after submitting his solution and during his problem reflection, Willibald states, *“Oh, I just noticed. I said there’ll be no effect on the interaction, but then I said van der Waals forces which is an interaction. So **I just contradicted myself in there**.”* After this recognition, Willibald decides to amend his first solution, ultimately improving his prediction. Similarly, when Sara was walking through her thought process during her problem reflection she noted, *“I guess I could **add onto my answer** that it could produce a van der Waals because of the close proximity.”* Importantly, Sara adds this correct idea to her final solution. We also observed one student, Jeffery, evaluating whether or not his solution answered the problem asked, which is notable because we also observed students evaluating in this way when solving problems in small groups (Halmo et al., 2022): *“I guess I can’t say for sure, but I’ll say this new amino acid form[s] a bond with the neighboring amino acids and results in a new protein shape. **The only issue with that answer is I feel like I’m not really answering the question**: Predict any new non-covalent interactions that might occur with such a mutation.”*

###### What happened after evaluating solution?

When students evaluated their solution, they either decided to stick with their original answer or amend their solution. Evaluating solution often resulted in students adding to or refining their final answer. However, these solution amendments were not always beneficial or in the correct direction because of limited content knowledge. In other work on the metacognition involved in changing answers, answer-changing neither reduced or significantly boosted performance (Stylianou-Georgiou & Papanastasiou, 2017). The fact that Mila’s evaluation of her solution led to a less correct answer whereas Willibald and Sara’s evaluation of their solutions led to more correct answers further contributes to the variable success of answer-changing on performance.

##### Evaluating Experience: Difficulty

Evaluating difficulty occurred when students assessed the difficulty level of the problem, whether it was difficult or easy for them. Kyle revealed his evaluation of difficulty after solving, when he said, *“**This one was a little more difficult for me**.”* He made this statement in reference to how determining the interactions valine could participate in was more challenging than determining the interactions serine could participate in during the Protein X problem. Students also compared the difficulty of the two problems we asked them to solve. For example, Elena determined that the Pathway Flux problem was easier for her compared to the Protein X problem in her problem reflection: *“**I didn’t find this question as hard** as the last question just cause it was a little bit more simple.”* In contrast, Elaine revealed that she found the Protein X problem challenging because of the open-ended nature of the question: *“**I just thought that was a little more difficult** because it’s just asking me to predict what possibly could happen instead of like something that’s like, definite, like I know the answer to.”*

###### What happened after evaluating difficulty?

When students assessed the difficulty level of the problems in this study, they usually evaluated the problems as difficult and not easy. They made this assessment of difficulty after solving.

##### Evaluating Experience: Feelings

Evaluating feelings occurred when students assessed how their emotions were connected to their thinking. For example, when making a prediction Clare acknowledged her intuition, *“I have a gut feeling that it [the mutation] would [affect the non-covalent interaction], but I don’t know why.”* Students exclusively revealed these emotions they were experiencing when they reflected on their thought process, which is why we consider them a part of evaluation. Interestingly though, the feelings they described were directly tied to their monitoring. We found that students associated negative emotions (nervousness, worry, and panic) with a lack of understanding or a lack of familiarity. For example, in Renee’s problem reflection, she connected feelings of panic to when she monitored a lack of understanding: *“I kind of **panicked** for a second, not really panicked cause I know this isn’t like graded or anything, but I do not know what a metabolic pathway is.”* In contrast, students associated more positive feelings when they reflected on moments of monitoring understanding or familiarity. For example, Renee stated, *“At first that was kind of **happy** because I knew what was going on.”* Additionally, some students revealed their use of a strategy explicitly to engender positive emotions or to avoid negative emotions, like Tabitha: *“I looked at the first box, I tried to break it up into certain sections, **so I did not get overwhelmed** by looking at it.”*

###### What happened after evaluating feelings?

When students reflected on the emotions connected with their thinking, they associated positive emotions with understanding, and negative emotions with not knowing or a lack of familiarity. Additionally, they identified the purpose of some strategy use to avoid negative emotions.

##### Implications for Instruction & Research about Evaluating

Our data indicate that some first-year life science students are evaluating their knowledge and experience during and after individual problem solving. As instructors, we can encourage students to evaluate their knowledge more by prompting them to 1) rethink or re-do a problem to see if they come up with the same answer or want to amend their first solution, and 2) predict if they think their solution is right or wrong. Encouraging students to evaluate by predicting if their solution is right or wrong is limited by the students’ individual content knowledge and accuracy. Therefore, it is imperative to help students develop their self-evaluation accuracy by following up their predictions with immediate feedback to help them become well-calibrated (Osterhage, 2021). Additionally, encouraging students to reflect on problem difficulty and the emotions involved with solving might help students identify and verbalize perceived barriers to problem solving to their instructors. There is likely a highly individualized level of desirable difficulty for each student where a problem is difficult enough to engage their curiosity and motivation to solve something unknown but also does not generate negative emotions associated with failure that could impede solving (de Bruin et al., 2023; Zepeda et al., 2020). The link between the emotional valence of feelings and metacognition in the present study is paralleled in other studies that used retrospective methods and found links between feelings of (dis)comfort and metacognition (Dye & Stanton, 2017). This suggests that the feelings students associate with their metacognition is an important consideration when designing future research studies and interventions. For example, helping students coach themselves through the negative emotions associated with not knowing and pivoting to what they do know might increase the positive emotions needed for problem-solving persistence.

### What aspects of learning self-efficacy do first-year life science students reveal when they solve problems on their own?

To address our second research question, we looked for statements related to self-efficacy in our participants’ think aloud data. Self-efficacy is defined as one’s belief in their capability to carry out a specific task (Bandura, 1997). Alternatively, self-efficacy is sometimes operationalized as one’s confidence in performing specific tasks (Ainscough et al., 2016). One motivational strategy that students use for increased self-efficacy is efficacy self-talk or “thoughts or subvocal statements aimed at influencing their efficacy for an ongoing academic task” (Wolters, 2003, p. 199). One form of efficacy self-talk that appeared in our data are self-encouraging statements we call “self-coaching”. These statements either 1) reassured themselves about a lack of understanding, 2) reassured themselves that it’s okay to be wrong, 3) encouraged themselves to keep going despite not knowing, or 4) reminded themselves of their prior experience. To highlight the role that self-coaching played in problem solving in our dataset, we first present examples where self-coaching was absent and could have been beneficial for the students in our study. Then we present examples where self-coaching was used.

#### When students monitored without self-coaching, they had hard time moving forward in their problem-solving

When solving the challenging biochemistry problems in this study, first-year life science students often came across pieces of information or parts of the figures that they were unfamiliar with or did not understand. In the Monitoring section, we described how students monitored their understanding and familiarity, but perhaps what is more interesting is *how students responded* to not knowing and their lack of familiarity. In a handful of cases, we witnessed students get stuck or hung up on what they did not know. We posit that the feeling of not knowing could increase anxiety, cause concern, and increase self-doubt, all of which can negatively impact a student’s self-efficacy and cause them to stop problem solving. One example of this in our data comes from Tiffani. Tiffani stated her lack of knowledge about how to proceed and followed this up with a statement on her lack of ability to solve the problem, *“I am actually not sure how to solve this. I do not think I can solve this one.”* A few lines later, Tiffani clarified where her lack of understanding rested, but again stated she cannot solve the problem, *“I’m not really sure how these type of amino acids pair up, so I can’t really solve it.”* In this instance, Tiffani’s lack of understanding is linked to a perceived inability to solve the problem.

Some students also linked not knowing with perceived deficits. For example, in the following quote Chandra linked not knowing how to answer the second part of the Protein X problem with the idea that she is *“not very good”* with non-covalent interactions: “*I’m not really sure about the second part. I do not know what to say at all for that, to predict any new non-covalent, I’m not very good with non-covalent at all.”* When asked where she got stuck during problem solving, Chandra stated, *“The “predict any new non-covalent” cause [I’m] not good with bonds. So I cannot predict anything really.”* In Chandra’s case, her lack of understanding was linked to a perceived deficit and inability to solve the problem. As instructors, it is moments like these where we would hope to intervene and help our students persist in problem solving. However, targeted coaching for all students each time they solve a problem can seem like an impossible feat to accomplish in large, lecture-style college classrooms. Therefore, from our data we suggest that encouraging students to self-coach themselves through these situations is one approach we could use to achieve this goal.

#### When students monitored and self-coached, they persisted in their problem-solving

In contrast to the cases of Tiffani and Chandra shared above, we found instances of students self-coaching after acknowledging their lack of understanding about parts of the problem by immediately reassuring themselves that it was okay to not know. For example, when exploring the arrows in the Pathway Flux problem figure Ivy states, *“I don’t really know what that little negative means, **but that’s okay**.”* After making this self-coaching statement Ivy moves on to thinking about the other arrows in the figure and what they mean to formulate an answer. In a similar vein, when some students were faced with their lack of understanding, one strategy they deployed was not dwelling on their lack of knowledge and pivoting to look for a foothold of something they do know. For example, in the following quote we see Viola acknowledge her initial lack of understanding and familiarity with the Pathway Flux problem and then find a foothold with the term enzymes which she knows she has learned about in the past, *“I’m thinking there’s very little here that I recognize or understand. Just… okay. So talking about enzymes, I know we learned a little bit about that.”*

Some students acknowledged this strategy of pivoting to what they do know in their problem reflections. In their problem reflections, Quinn and Gerald expanded that they will rely on what they do know, even if it is not accurate. As Quinn put it, *“**taking what I think I know, even if it’s wrong**, like I kind of have to, you have to go off of something.”* Similarly, Gerald acknowledged his strategy of *“**it’s okay to get it wrong**”* when he doesn’t know and connects this strategy to his experience solving problems on high-stakes exams,

> *I try to use information that I knew and I didn’t know a lot. So I had to kind of use my strategy where I’m like, if this was on a test, this is one of the questions that I would either skip and come back to or write down a really quick answer and then come back to**. So my strategy for this one is it’s okay to get it wrong. You need to move on and make estimated guess.** Like if I wasn’t sure what the arrows meant, so I was like, ”okay, make an estimated guess on what you think the arrows mean. And then using the information that you kind of came up with try to get a right answer using that and like, explain your answer so maybe they’ll give you half points…” – Gerald*

We also observed students encouraging themselves to persist despite not knowing. In the following quote we see Kyle acknowledge a term he doesn’t know at the start of his think aloud and verbally choose to keep going, *“So the title is pathway flux problem. I’m not too sure what flux means, **but I’m going to keep on going**.”* Sometimes this took the form of persisting to write an answer to the problem despite not knowing. For example, after Kathleen states, *“I don’t know what flux is. That’s okay.”* she goes on to say, *“Through the pathway as whole, probably. Okay. **I’m going to try and answer it now**.”* Additionally, take Viola’s statement of, *“I’m not even really sure what pathway flux is. So I’m also not really sure what the little negative sign is and it pointing to B. **But I’m going to try to type an answer**.”* Rather than getting stuck on not knowing what the negative feedback loop symbol depicts, she moves past it to come to a solution.

We also saw students use self-coaching to remind themselves of their prior experience. In the following example, we see Mila talk herself through the substitution of serine with valine in the Protein X problem: *“So there’s not going to be a hydroxyl anymore, but I don’t know if that even matters, but there, valine, has more to it. I don’t know if that means there would be an effect on the covalent interaction. I haven’t had chemistry in such a long time [pause], but at the same time, **this is bio. So I should still know it.** [laughs]”* Mila’s tone as she made this statement was very matter-of-fact. Her laugh at the end suggests she did not take what she said too seriously. After making this self-coaching statement, Mila rereads the question a few times and ultimately decides that the non-covalent interaction is affected because of the structural difference in valine and serine. Prior experiences, sometimes called mastery experiences, are one established source of self-efficacy that Mila might have been drawing on when she made this self-coaching statement (Bandura, 1977; Pajares, 1996).

##### Implications for Instruction about Self-Coaching

Students can be encouraged to self-coach by using some of the phrases we identified in our data as prompts. However, we would encourage instructors to rephrase some of self-coaching statements in our data by removing the word “should” because this term might make students feel inadequate if they think they are expected to know things they don’t yet know. Instead, we could encourage students to remind themselves of when they’ve successfully solved challenging biology problems in the past by saying things like, “I’ve solved challenging problems like this before, so I can solve this one.” Taken together, we posit that self-coaching could be used by students to decrease anxiety and increase confidence when faced with the feeling of not knowing that can result from monitoring, which could potentially positively impact a student’s self-efficacy and metacognitive regulation. Our results reveal first-year students are monitoring in a myriad of ways. Sometimes when students monitor, they may not act further on the resulting information because it makes them feel bad or uncomfortable. Self-coaching could support students to act on their metacognition or not actively avoid being metacognitive.

## LIMITATIONS

Even with the use of in-the-moment methods like think aloud interviews, we are limited to the metacognition that students verbalized. For example, students may have been employing metacognition while solving that they simply did not verbalize. However, using a think aloud approach in this study ensured we were accessing students’ metacognition in use, rather than their remembrance of metacognition they used in the past which is subject to recall bias (Schellings et al., 2013). Our study, like most education research, may suffer from selection bias where the students who volunteer represent a biased sample (Collins, 2017). To address this potential pitfall, we attempted to ensure our sample represented the student body at each institution by using purposeful sampling based on demographics and varied responses to the revised Metacognitive Awareness Inventory (Harrison & Vallin, 2018). Lastly, while our sample size is large (N = 52) for qualitative analyses and includes students from three different institutional types, the data are not necessarily generalizable to contexts beyond the scope of the study.

## CONCLUSION

The goal of this study was to investigate first-year life science students’ metacognition and self-efficacy in-the-moment while they solved challenging problems. Think aloud interviews with 52 students across three institutions revealed that first-year life science students use an array of monitoring and evaluating skills while solving challenging problems but put less emphasis on planning. We also found instances of students self-coaching or encouraging themselves when confronted with a lack of understanding or a lack of familiarity, which helped them use their metacognition to take action and persist in problem solving. Oftentimes, researchers studying metacognition can find themselves unintentionally operating from a deficit standpoint. However, our findings challenge the notion that first-year life science students enter college with poorly developed metacognitive skills. Indeed, the first-year life science students in this study were monitoring and evaluating when solving challenging biology problems on their own. Together these findings about in-the-moment metacognition and self-efficacy offer a positive outlook on ways we can encourage students to couple their developing metacognitive regulation skills and self-efficacy to persist when faced with challenging disciplinary problems.

## ACKNOWLEDGMENTS

We would like to thank Dr. Paula Lemons for allowing us to use problems developed in her research program for this study and for her helpful feedback during the writing process, the College Learning Study participants for their willingness to participate in this study, Dr. Mariel Pfeifer for her assistance conducting interviews and continued discussion of the data during the writing of this manuscript, C.J. Zajic for his contribution to preliminary data analysis, and Emily K. Bremers, Rayna Carter, and the UGA BERG community for their thoughtful feedback on earlier versions of this manuscript. This material is based on work supported by the National Science Foundation under Grant Number 1942318. Any opinions, findings, and conclusions or recommendations expressed in this material are those of the authors and do not necessarily reflect the views of the National Science Foundation.

