## Supplemental Materials for "Metacognition and Problem Solving: How Self-Coaching Helps First-Year Students Move Past the Discomfort of Monitoring"

**Supplemental Materials**
**Table of Contents**

**Think Aloud Interview Protocol………………………………………………………………………….2**

**Supplemental Data……………………………………………………………………………………6**

**Problem-Solving Vignettes…………………………………………………………………………….7**

**Think Aloud Interview Protocol**

### Think-Aloud Description

*"In this part of today's interview, I'll ask you to think aloud while you work through two biology problems. Think aloud means that you say out loud whatever comes into your mind as you work through problems and as you write down your answer. Please say anything that comes into your mind, even, "I don't know what to say next." I will also remind you to think aloud or I might ask you to say more about what you've said in order to understand more about what you are thinking. We'll practice first with a generic problem. Before we begin with the practice problem, what questions do you have about this part of the interview?"*

### Practice Problem

*"If you're ready, let's practice. Please remember to say everything that comes into your mind as you work through this problem."* [use the share screen feature in Zoom to share page 1 of the "Year 1 Problem Set.pdf" file]

**Practice Problem**

While growing vegetables in your backyard, you noticed a particular kind of insect eating your plants. You took a rough count (see data below) of the insect population over time.

| Time (days) | Insect Population (number) |
|---|---|
| 2 | 7 |
| 4 | 16 |
| 8 | 60 |
| 10 | 123 |

Which graph shows the best representation of your data?

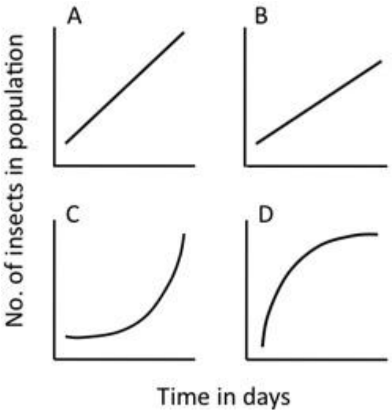


Use the following prompts as necessary:

- *"What are you thinking (now)?"*
- *"Can you tell me more about that?"*

*"Please type your solution to the Practice Problem in the Zoom chat box."* [wait for participant to type]

**Practice Problem Reflection**

*"Thank you. Now that you have solved the practice problem, please describe how you solved this problem. List the steps of your own thinking process, in order from start to finish. Provide as much detail as you can."*

**Biology Problems**

*"Great. Now that you've had a chance to practice, I'll give you the biology problems. These biology problems may be challenging, and that is okay. Coming to a correct solution is not the goal. Do the best you can to solve the problems and write your answers in the Zoom chat box. We are not grading you. Rather, we are genuinely interested in how you approach these challenging problems. For this reason, we ask that you do not use outside resources while working through these problems. You may take as long as you need, but we will probably be done with this part in about 20 or 30 minutes. Please remember to say everything that comes into your mind while you are solving and writing an answer to the problems. Let's begin."* [scroll down to page 2 of the "Year 1 Problem Set.pdf" file]

**Protein X Problem**
**(Figure 3 from Halmo et al., 2020)**

Protein X, a cytoplasmic protein, is folded into its tertiary structure, surrounded by water molecules (red and gray). This environment has a pH of 7.4. The blue line represents the protein X backbone. Some, but not all, of the amino acid side chains are shown in chemical notation.

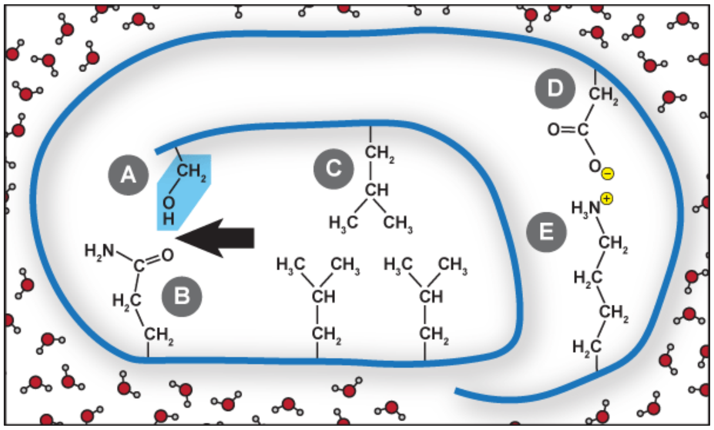


The amino acids shown are: (A) serine, (B) glutamine, (C) leucine, (D) aspartate, and (E) lysine. Sometimes, a mutation occurs that substitutes serine (blue highlight) with valine (below).

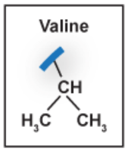


Do you predict that such a mutation would affect the non-covalent interaction pointed to by the arrow? Predict any new non-covalent interactions that might occur with such a mutation, and provide a scientific explanation to support your prediction.

Use the following prompts as necessary:

- *"What are you thinking (now)?"*
- *"Can you tell me more about that?"*

*"Please type your solution to the Protein X Problem in the Zoom chat box."* [wait for participant to type]

**Protein X Problem Reflection**

*"Thank you. Now that you have solved the first problem, please describe how you solved this protein X problem. List the steps of your own thinking process, in order from start to finish. Provide as much detail as you can."*

Use the following prompts as necessary:

- *What did you think the problem was asking?*
- *What did you already know or remember when solving?*
- *What strategies did you use to solve?*
- *If you got stuck, where did you get stuck, and what do you wish you could have looked up?*

*"Thank you for sharing your thoughts while you solved the first problem. Let's move on to the next one."* [scroll down to page 3 of the "Year 1 Problem Set.pdf" file]

**Pathway Flux Problem**
**(Figure 1 from Bhatia et al., 2022)**

The metabolic pathway below shows the conversion of thiamine phosphate to dimethylacryloyl-CoA (DMA-CoA).

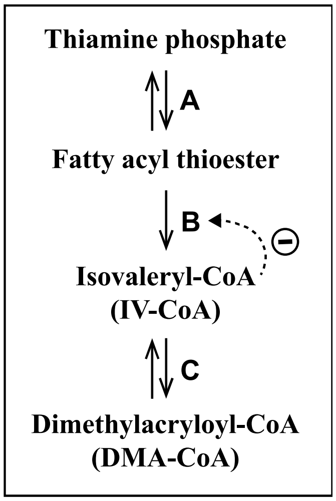


A change occurs so that IV-CoA can no longer bind to enzyme B. Do you predict that this will affect flux through the pathway as a whole? Provide a scientific explanation to support your prediction.

Use the following prompts as necessary:

- *"What are you thinking (now)?"*
- *"Can you tell me more about that?"*

*"Please type your solution to the Pathway Flux Problem in the Zoom chat box."* [wait for participant to type]

**Pathway Flux Problem Reflection**

*"Thank you. Now that you have solved the second problem, please describe how you solved this pathway flux problem. List the steps of your own thinking process, <u>in order from start to finish</u>. Provide as much detail as you can."*

Use the following prompts as necessary:

- *What did you think the problem was asking?*
- *What did you already know or remember when solving?*
- *What strategies did you use to solve?*
- *If you got stuck, where did you get stuck, and what do you wish you could have looked up?*

[stop sharing screen]

**Closing Questions**

*"Thank you for solving those challenging biology problems. These problems are from a biochemistry class, so if the problems seemed unfamiliar that is okay. You did a good job working through them by drawing upon your prior knowledge from biology and chemistry."*

1) On a scale from 1-10, with 10 being the highest, how would you rate your level of understanding of noncovalent interactions and metabolism?

2) What challenges, if any, did you experience when solving the problems today?

3) What would you do to obtain a deeper understanding of noncovalent interactions and metabolism?

### Supplemental Data

**Supplemental Table 1. Comparison of Data Collection Sites**

| | Georgia Gwinnett College | University of Georgia | University of North Georgia |
|---|---|---|---|
| Institution Type | Baccalaureate College | Doctoral R1 | Master's University |
| Setting | Suburban | City | Suburban |
| Number of undergraduates | 10,949 | 28,848 | 18,173 |
| Students from racially minoritized groups | 58.1% | 14.1% | 24.0% |
| Students who identify as women | 58.7% | 57.7% | 56.2% |
| Students who identify as first-generation | 40% | 7.65% | 21.5% |
| Average High School GPA | 2.9 | 4.0 | 3.6 |
| Average SAT Score | 1012 | 1281 | 1108 |

**Supplemental Table 2. Participant Demographics by Institution**

| | Georgia Gwinnett College | University of Georgia | University of North Georgia |
|---|---|---|---|
| Number of participants | 8 | 23 | 21 |
| Participants from racially minoritized groups | 4 | 5 | 5 |
| Participants who identify as women | 8 | 13 | 15 |
| Participants who identify as first-generation | 5 | 3 | 6 |
| Average High School GPA | 3.3 | 4.0 | 3.6 |
| Average College GPA | 3.4 | 3.7 | 2.9 |

*Note: For average high school GPA, institutional data is missing from two GGC students.*

**Supplemental Table 3. Problem-Solving Performance by Institution**

| Institution | Mean Problem-Solving Performance | Standard Deviation |
|---|---|---|
| GGC | 1.88 | 0.64 |
| UGA | 1.78 | 0.62 |
| UNG | 1.86 | 0.80 |

**Supplemental Table 4. Problem-Solving Performance by Gender**

| Gender | Mean Problem-Solving Performance | Standard Deviation |
|---|---|---|
| Man | 1.69 | 0.81 |
| Woman | 1.89 | 0.62 |

### Problem-Solving Vignettes

The three following vignettes represent more detailed snapshots of the metacognitive regulation we observed during individual problem-solving of the Protein X problem. We provide a vignette from a single student from each institution. For each individual student, we offer areas in which they each could further develop their metacognitive regulation skills since metacognitive interventions likely need to be tailored to the individual and context.

#### Ignacio from UGA

Even though Ignacio didn't come to a correct answer for the Protein X problem, he displayed the use of monitoring and evaluating skills in his think aloud.

Ignacio's path to a solution involved cycles of monitoring and evaluating. Ignacio starts his think aloud by reading the problem. As he reads, Ignacio **monitors which given information in the problem stem might be relevant**: "This environment has a pH of 7.4. So maybe the pH has something to do with it." Monitoring relevance is a large part of Ignacio's process. After reading the problem, he **assesses the task** and mentally places valine where the serine was in the figure: "Okay. So it says serine is substituted with valine. So now that I understand that, place valine there…" After making this mental substitution, he continues to **monitor what relevant parts of the problem he should focus on**. In deciding to focus on valine, he reveals an inaccurate idea that serine is not interacting with anything around it: "So, first things first, it doesn't even matter, serine isn't having really any interactions with any surrounding chemicals. So it all comes down to valine." Ignacio points out relevant functional groups within the glutamine amino acid ("carbon formed two bonds with an oxygen over there") and the valine amino acid ("CH3") and goes on to comment of the **irrelevance of the other provided information**: "So the pH has nothing to do with it. The water molecules had nothing to do with it as well. So basically everything in that first half, everything in that first thing, right there is basically useless. So I'm just going to exclude that information out of my thought process cause the pH has nothing to do with what's going on right now and the glutamine, leucine, aspartate, lysine. So all you have to worry about is glutamine because it seems like (A) only has the opportunity based on distance in the graph to interact with glutamine or leucine. (D) and (E) are out." This monitoring was likely helpful for Ignacio as he solved, however we note that the pH is relevant information for determining the ionization state of certain amino acids and that water plays an important role in protein folding. Having narrowed down his focus on the glutamine and valine, Ignacio predicts that a non-covalent interaction could form between these two amino acids, specifically between a carbon atom in valine and the oxygen atom in glutamine. However, Ignacio does not name this interaction or describe how it would form. As he is writing his answer, he starts thinking about the difference between covalent bonds and non-covalent bonds which prompts him to **evaluate the accuracy of his solution** and then **correct** his definition in the wrong direction, "I'm trying to think. Covalent bonds versus non-covalent bonds. I think I'm right in that justification cause a covalent bond would build up a sound… no, covalent bonds are between, so hydrogen bonds are covalent bonds, hmm so non-covalent interactions between the carbon atom and H two C." Ignacio's self-correction is initially in the wrong direction because hydrogen bonds are actually a type of non-covalent interaction between molecules and he describes the covalent bonds within the molecule of valine as non-covalent. He ultimately **recorrects** in the right direction when he states, "So yeah, I think that would form a non-covalent interaction, cause it wouldn't be a strong bond because it wouldn't be the same as covalent, which is holding that, the valine together. It just may interact together…"

As Ignacio submits his final solution he **assesses the accuracy of his answer and expresses tentativeness about his understanding of covalent and non-covalent** because he hasn't reviewed that material recently, "Okay. So I feel right on that. And yeah, so then that's, that's my answer, but again, I haven’t really gone over covalent and non-covalent bonds in a minute, so I'm just hoping that's something that makes sense.” In his reflection, Ignacio **acknowledges the assumption he made about covalent versus non-covalent**, “And then getting to think about what covalent and non-covalent bonds are. I don't know if I'm correct in this assumption, but I was assuming covalent is the bonds between the ions inside of each amino acids. Like the carbon and oxygen have a covalent bond, but non-covalent bonds could occur between ions that aren't bound in the same amino acid.” He also **evaluates the difficulty of the problem** by noting that the problem would have been simple if he could check his understanding of the definitions of these terms: “...if I had a greater understanding of definitions of covalent and non-covalent, I think the question would be quite simple, but due to the fact that I have not gone over non-covalent and covalent bonds, about four years since AP bio I'm guesstimating on what my initial knowledge was.” Ignacio goes on to identify this gap in knowledge as what he wished he could have looked up when solving. In the closing section of the interview when describing his self-assessment of understanding he noted, “But the thought process that I developed over the years of science-based learning has led to me kind of like, you know, thinking in a way that led me to be able to do probably what they meant based off of my own understanding… the thought processes that apply to every science class led to it kind of being kind of a little, or made me feel more confident, probably than I should’ve.” Through this statement it is clear that Ignacio is aware of his confidence and how his high confidence levels could be potentially misleading. **Implications for Instruction: Students like Ignacio could benefit from practice and feedback on calibrating their monitoring of their understanding to actual performance.**

**Harper from GGC**

Despite not coming to a correct answer for the Protein X problem, Harper demonstrated awareness of the limits of her knowledge and used monitoring skills in her think aloud.

Harper’s path to a solution involved several types of monitoring. First, Harper read the Protein X problem out loud, then immediately reread the question to verify what it was asking. She focused her attention on the arrow, then reread the problem again. Harper **monitored her understanding** by stating what she knew about the term covalent, “And now I'm thinking of the definition of covalent. It would be bonds sharing equally or not equally…” After monitoring her understanding of the term covalent in this way, Harper **revealed her confusion and lack of knowledge about how to proceed and ties this to her uncertainty about monitoring the relevance of the provided information**: “Hmm. I'm not sure what to do next. I'm not sure what to think next… I'm not sure how to approach the question. Like, I'm not sure If I'm comprehending the information that was given. If the information was necessary.” After stating this, she focused on the blue highlighted area, started looking at the molecules, and then **reread the question as a strategy to employ when she is confused**: “Now I'm just rereading the question, cause I don't think I much understand what I'm supposed to be looking at.” After employing this strategy, Harper **accurately assesses the task** by “thinking about what it would look like if valine were in the place of the blue highlighted substance”. Harper then makes a tentative prediction and notes her **lack of knowledge about how to proceed** because **she is uncertain about her ability to answer due to a lack of prior knowledge** : “I think that I wanna say that if valine were swapped out with serine then it would affect the non-covalent interaction, but I'm not sure if I'd be able to answer the second question of the question, the predict any new non-covalent interactions that might occur…I don't think that I have enough like basis or learning to where I'm able to answer that question. I'm not sure if I'd be able to predict seeing as I don't know much about the topic or much about how to answer this question.” In her reflection, Harper revealed that the second part of the question was where she got stuck, which is an accurate assessment. However, Harper said she would look up “what a mutation would do to a non-covalent interaction.” While this might have helped Harper in her problem solving, she may have benefitted more from looking up the different types of non-covalent interactions first. **Implications for Instruction: Students like Harper could benefit from learning more about effective strategies they can use when they experience feeling stuck during problem solving.**

**Sara from UNG**

Despite not coming to a complete and fully correct answer for the Protein X problem, Sara was the highest performer in our sample. In her think aloud, she looped through cycles of monitoring correctness to generate and revise her prediction and evaluated her solution as she reflected.

Sara's path to a solution involved cycles of monitoring and predicting followed by evaluating her final solution. To start, Sara reads the Protein X problem once through and **identifies that the current interaction in question is a hydrogen bond**, "So non-covalent, so it can just bond, and then that'd be hydrogen oxygen. So that'd be hydrogen bond, because they are kind of just chilling there, and then. But that also has hydrogen." In the same fell swoop she also **notices that the mutation, valine, also contains hydrogens**. Sara goes on to **incorrectly predict** that with the mutation, "It would still have the capabilities of having hydrogen bond because of the different charges." However, she immediately **monitors the correctness of her prediction and modifies it**: "So. No. Maybe it'd be stronger. It could be stronger because it has no, no, it, it would have a change." Sara's correction is initially in the wrong direction. The interaction between glutamine and valine would not be stronger than the interaction between glutamine and serine. However, she **monitors the correctness of her prediction again** and decides there would be a change in the interaction. After making this prediction to answer the first part of the problem, Sara **rereads** the second part of the problem. She then goes on to **predict new non-covalent interactions by going back to her original prediction** that the hydrogen bond could still form but considers the opposite idea from before, that the interaction wouldn't be as strong because valine has more hydrogens than serine: "It would be about the same, but then it could also bond to more things. So I guess. So at the heart, but there is three hydrogens. So there's only room for one in the hydrogen bond. And so it'd still be a hydrogen bond, I guess it could bond to more things. Maybe, bond wouldn't be strong?" However, Sara stops her train of thought and **monitors the correctness of this incorrect prediction once again** when she says, "Wait. No, no. Cause the O and the H have different charges. That's a covalent bond between carbon and hydrogen, and covalent bonds don't have, oh, then yeah. I wouldn't be able to bond. So it yeah, it wouldn't be able to bond anymore. No, I think, yeah, I'm going to go with that…" Through monitoring the correctness of her ideas, Sara comes to the correct idea that the oxygen and hydrogen have different charges and that with the substitution of valine for serine, the hydrogen bond would no longer be able to form.

During the reflection after she provided her solution, Sara reveals that she **monitored the relevance of the provided information** and determined that it was irrelevant for solving the problem: "So at first I read over the thing, and then there was a lot of unneeded information in the first little paragraph. I mean, it told you what it was and like the pH and stuff, but you really didn't need it to understand the bonding. Then I, when I read the amino acid shown, I just kind of kept that information aside. Cause you don't really need to know that in order to predict the interactions and stuff, all that you really need to know is that when oxygen and hydrogen bonds, they leave partial charges." Sara also **continues to reveal more of her understanding when she classifies the charges she mentioned earlier as "partial"**. She struggled to identify the exact partial charges and while **monitoring the correctness of the charges**, she gets exasperated, "So they don't really have, they don't have the charges that oxygen and hydrogen have. So you don't have the negative partial and the negative, negative partial, negative, positive and negative. Wait, no partial positive. Partial negative. Oh good grief - to create the hydrogen, but I guess, I mean, I guess I could add onto my answer that it could produce a van der Waals because of the close proximity." Sara doesn't get too hung up on the exact partial charges and **evaluates her solution** by identifying she could add more to her answer about the van der Waals interactions that would happen between the valine and glutamine. It is notable that Sara revealed even more of her metacognition in the reflection portion of the think aloud that she did not share verbally as she was concurrently solving. **Implications for Instruction: Students like Sara might have automatic metacognitive regulation skills who could benefit from metacognitive prompting alone. For example, Sara's initial solution could have been more complete and correct if she had been prompted to reflect on her solution.**
